# Comparative transcriptomics reveal a highly polymorphic *Xanthomonas* HrpG virulence regulon

**DOI:** 10.1101/2024.05.17.594730

**Authors:** Thomas Quiroz Monnens, Brice Roux, Sébastien Cunnac, Erika Charbit, Sébastien Carrère, Emmanuelle Lauber, Marie-Françoise Jardinaud, Armelle Darrasse, Matthieu Arlat, Boris Szurek, Olivier Pruvost, Marie-Agnès Jacques, Lionel Gagnevin, Ralf Koebnik, Laurent D. Noël, Alice Boulanger

**Affiliations:** LIPME, Université de Toulouse, INRAE/CNRS/Université Paul Sabatier Toulouse 3, UMR 0441/2598, Castanet-Tolosan, France; Univ. Angers, Institut Agro, INRAE, IRHS, SFR QUASAV, F-49000 Angers, France; CIRAD, UMR PVBMT, F-97410 Saint-Pierre, La Réunion, France; PHIM, Université de Montpellier, IRD, CIRAD, INRAE, Institut Agro, Montpellier, France; CIRAD, UMR PHIM, F-34398 Montpellier, France

**Keywords:** *Xanthomonas*, HrpG, type III secretion system, regulon diversity, transcriptome sequencing

## Abstract

Bacteria of the genus *Xanthomonas* cause economically significant diseases in various crops. Their virulence is dependent on the translocation of type III effectors (T3Es) into plant cells by the type III secretion system (T3SS), a process regulated by the master response regulator HrpG. Although HrpG has been studied for over two decades, its regulon across diverse *Xanthomonas* species, particularly beyond type III secretion, remains understudied. In this study, we conducted transcriptome sequencing to explore the HrpG regulons of 17 *Xanthomonas* strains, encompassing six species and nine pathovars, each exhibiting distinct host and tissue specificities. We employed constitutive expression of plasmid-borne *hrpG*\*, which encodes a constitutively active form of HrpG, to induce the regulon. Our findings reveal substantial inter- and intra-specific diversity in the HrpG* regulons across the strains. Besides 21 genes directly involved in the biosynthesis of the T3SS, the core HrpG* regulon is limited to only five additional genes encoding the transcriptional activator HrpX, the two T3E proteins XopR and XopL, a major facility superfamily (MFS) transporter, and the phosphatase PhoC. Interestingly, genes involved in chemotaxis and genes encoding enzymes with carbohydrate-active and proteolytic activities are variably regulated by HrpG*. The diversity in the HrpG* regulon suggests that HrpG-dependent virulence in *Xanthomonas* might be achieved through several distinct strain-specific strategies, potentially reflecting adaptation to diverse ecological niches. These findings enhance our understanding of the complex role of HrpG in regulating various virulence and adaptive pathways, extending beyond T3Es and the T3SS.

**IMPORTANCE:** In the decades since its discovery, HrpG and its role in the regulation of the type III secretion system (T3SS) and its associated type III effectors (T3Es) in *Xanthomonas* has been the subject of extensive research. Despite notable progress in understanding its molecular regulatory mechanisms, the full spectrum of processes under control of HrpG, particularly beyond the T3SS and T3Es, and the degree of regulatory conservation across plant-pathogenic *Xanthomonas* species, remained unclear. To address this knowledge gap, we systematically compared the transcriptomes of 17 *Xanthomonas* strains, expressing a constitutively active form of HrpG, called HrpG*. We showed that HrpG* regulates different physiological processes other than the T3SS and T3Es and that this regulation shows substantial variation across the different strains. Taken together, our results provide new insights into *Xanthomonas-*plant interactions through the regulation of different metabolic and virulence pathways by the master response regulator HrpG.

## INTRODUCTION

In phytopathogenic *Xanthomonas* bacteria, HrpG is a conserved response regulator belonging to the OmpR family of two-component regulatory systems (1). Two-component regulatory systems are one of the major mechanisms used by bacteria to perceive and adapt their physiology to changing environmental conditions (2). These systems typically rely on a phosphorelay between a sensor kinase and a response regulator, which becomes active upon phosphorylation. In *Xanthomonas*, HrpG regulates the expression of *hrpX,* which encodes a transcriptional regulator. HrpX binds to the plant-inducible promoter (PIP) box (3), a DNA sequence present in the cis-elements of various pathogenicity genes, including type III effector proteins (T3Es) and the *hrp* (hypersensitive response and pathogenicity) genes, which encode the type III secretion system (T3SS) (4, 5). This system is one of the main virulence factors of *Xanthomonas* and it enables the direct delivery of bacterial T3Es into the plant cellular environment, where they interfere with plant immunity and alter plant physiology to facilitate disease (6). Disruption of any key structural component of the T3SS results in a complete loss of pathogenicity for *Xanthomonas* species (4).

The *Xanthomonas* genus comprises 33 species of Gram-negative □-proteobacteria (7) causing diseases on more than 400 different monocot and dicot plants including economically-important crops (8). *Xanthomonas* species are further divided into pathovars that can exhibit distinct host specificities and tissue tropisms, causing a variety of symptoms such as wilting, necrosis or blight (8). Phylogenetic group-I *Xanthomonas* include monocot pathogens such as *X. translucens* pv. *translucens* (*Xtt*) (9), causal agent of bacterial leaf streak of barley. On the other hand, phylogenetic group-II *Xanthomonas* can infect both monocot and dicot plants. For example, *X. oryzae* pv. *oryzae* (*Xoo*) causes bacterial blight on rice, a monocot, while *X. phaseoli* pv. *phaseoli* (*Xpp*) and *X. citri* pv. *fuscans* (*Xcf*) belonging to phylogenetically distinct species are both able to cause common blight of common bean, a dicot. Other economically important *Xanthomonas* pathovars include *X. citri* pv. *mangiferaeindicae* (*Xcm*), causal agent of mango bacterial black spot, *X. citri* pv. *citri* (*Xci*), responsible for Asiatic citrus canker and *X. euvesicatoria* pv. *alfalfae* (*Xea*), which causes disease on legumes. The genus also comprises *X. campestris* pv. *campestris* (*Xcc*) and *X. campestris* pv. *raphani* (*Xcr*), the causal agents of black rot disease and bacterial spot disease of Brassicaceae respectively. *Xcc*, *Xoo* and *Xtt* are known to infect the plant vasculature through wounds or hydathodes while *Xci*, *Xcm*, *Xea* and *Xcr* colonize the mesophyll through stomata. *Xcf* and *Xpp* colonize their host plant by both ways (10–13).

Since its discovery, research has shown that HrpG regulates a broad array of genes beyond T3Es and the T3SS. This broader regulatory role was initially identified by comparing the transcriptomes of wild-type versus *hrpG* mutant strains in *hrp*-inducing media (14) or wild-type strains versus strains expressing an auto-active gain-of-function form of HrpG, such as HrpG^E44K^, commonly named HrpG* (1, 15, 16). Additional studies have supported these claims, showing HrpG could regulate the expression of genes other than those associated with type III secretion, including genes associated with chemotaxis and motility, and genes encoding extracellular proteases and cell wall degrading enzymes associated with the Xps type II secretion system (T2SS) (17–20). However, this part of the HrpG regulon remains understudied and appears to be poorly conserved between *Xanthomonas* species.

To investigate the conservation of the HrpG regulon across *Xanthomonas* species, we performed a comprehensive transcriptomic and genomic analysis of 17 *Xanthomonas* strains transiently expressing the constitutively active *hrpG\** variant (21). The HrpG* regulons identified in the different strains varied considerably in their size, ranging from 137 to 2,355 genes. Interestingly, the core HrpG* regulon across these 17 strains comprises only 26 genes, mainly involved in the biogenesis of the T3SS itself. Moreover, it was found that the full extent of genes and processes regulated by HrpG* is remarkably diverse across the 17 strains. These findings suggest that HrpG-dependent pathogenicity in *Xanthomonas* species can be achieved through diverse strategies.

## RESULTS

### Transcriptome-based structural annotation of 17 *Xanthomonas* genomes

17 *Xanthomonas* strains belonging to nine pathovars were selected to study the diversity and conservation of the HrpG regulon (Fig. 1A). These strains represent pathogens associated with diverse host plants, were sampled over nearly a hundred years in ten countries, and have various lifestyles (Table S1A; Fig. 1B). Genome sequences of strains *Xtt*_CFBP2054_ *,Xcc*_CN05_, *Xcm*_LG56-10_ and *Xcm*_LG81-27_ were determined using either short- (Illumina) or long-read (PacBio) sequencing (Table S1B; S1C). High-quality genome sequences were readily available for the 13 other strains.

**Figure 1.**
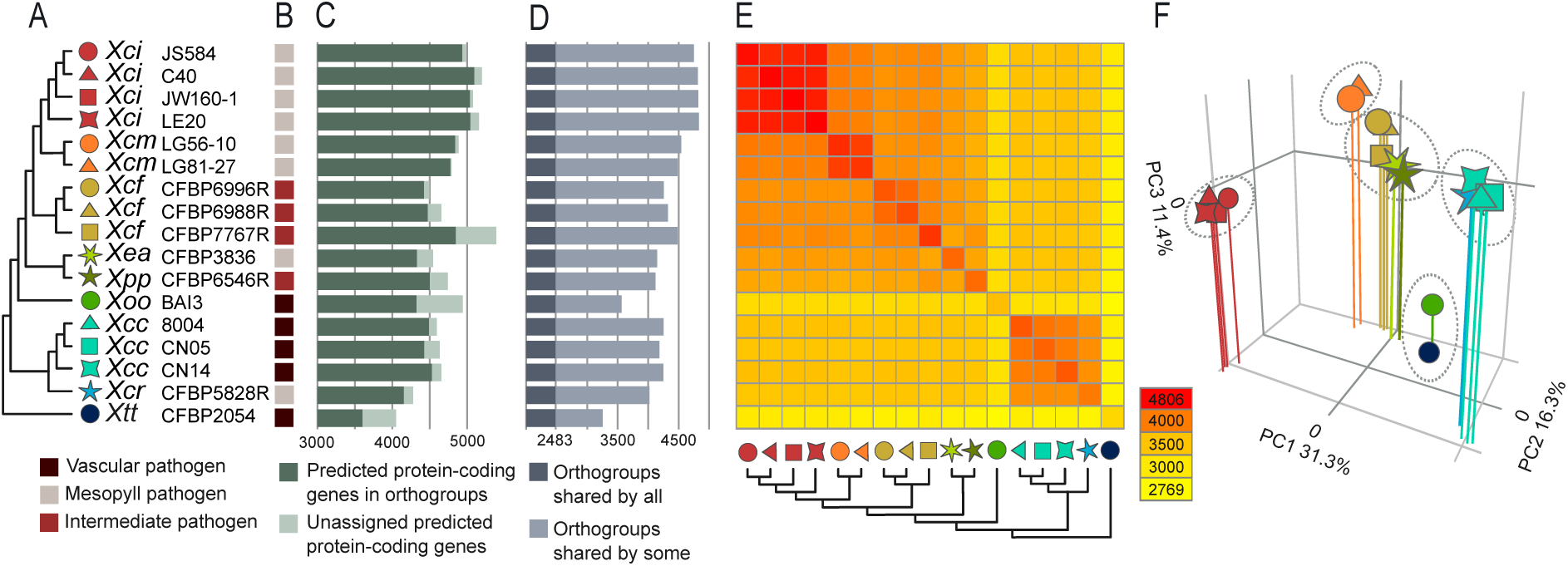
Gene orthology analysis of the 17 *Xanthomonas* strains used in this study. **A:** phylogenetic tree of all strains, each represented with a distinct coloured symbol. **B:** Lifestyle indicates if the pathogen infects primarily leaf mesophyll (light brown), the vascular tissues (dark brown) or both (burgundy). **C:** Number of predicted protein-coding genes per genome, including those in orthogroups (dark green) and singletons (light green). **D:** Number of orthogroups per genome. **E:** Heatmap showing the number of shared orthogroups between strains. Symbols as in panel **A**. **F:** Principal component (PC) analysis plot showing the clustering of the different strains based on the presence or absence of orthogroups against the first three PCs. Shape and color of symbols identifies each of the 17 strains from the nine pathovars studied as in panel **A**.

Experimentally-based and homogeneous annotation of genomes is vital for cross-species comparative transcriptomic analyses. Such annotations were previously made for *Xcc*_8004_ and *Xcr*_CFBP5828R_, which were based on various transcriptomic datasets and the EuGene-PP pipeline (16, 22). The same strategy was used to annotate the remaining genomes in combination with different RNA-Seq libraries (Table S1E), including those from conjugates carrying an empty vector or a vector harboring *hrpG\**. For *Xci* and *Xcm* strains, empty vector conjugates were not generated, and RNA-Seq libraries from wild-type strains were used for annotation instead. Small RNA sequencing was performed for several strains in order to identify small RNAs and determine transcriptional start sites. The newly annotated genomes consist of 4066 (*Xtt*_CFBP2054_) to 5421 (*Xcf*_CFBP7767_) protein-coding genes (Table S1C; Data S1 https://doi.org/10.57745/9OTYNJ). The 17 homogeneously annotated genomes were used for all downstream analyses.

### The *Xanthomonas* core genome is composed of 2,483 orthogroups

To enable cross species comparative transcriptomics, an orthogroup (OG) database was built with the newly annotated genomes using Orthofinder. Collectively, the 17 genomes contained 88,480 genes, of which 81,490 were predicted to be protein-coding. Out of these, 78,225 were assigned to a total of 6,899 orthogroups (Fig. 1C). 2,483 orthogroups were represented by at least one ortholog in each strain, representing the *Xanthomonas* core genome in this study (Fig. 1D; Data S2 https://doi.org/10.57745/9OTYNJ). A phylogenetic tree including all strains was built using the STAG algorithm and rooted using STRIDE (Fig. 1A). As expected, group-I *Xanthomonas* strain *Xtt*_CFBP2054_ was located at the root of the tree. To assess the overall similarity in orthogroup content among the strains, a heatmap depicting the number of shared orthogroups and a principal component analysis (PCA) based on a binary presence/absence matrix of all orthogroups in the dataset were used (Fig. 1EF). The high number of shared orthogroups between related strains in the heatmap, and the clustering of related strains in the PCA is consistent with the genetic relationships of the *Xanthomonas* strains.

### The accessory HrpG* regulons are highly diverse

The HrpG* regulons were identified by sequencing cDNAs of either wild-type strains with or without an empty plasmid or a plasmid carrying *hrpG\**. The biological reproducibility of each triplicate per genotype was evaluated through MultiQCs on mapping statistics, and through PCA, MA (M = log fold change, A = mean of normalized counts) and volcano plots of the DESeq2 output (Data S3, https://doi.org/10.57745/9OTYNJ). The reproducibility across all samples was consistent except for one *Xtt* replicate, which was excluded from subsequent differential expression analysis. As expected for a direct HrpG target, expression of the *hrpX* gene was significantly upregulated in all *hrpG*\* samples (Fig. 2AB). This demonstrates the presence of biologically active HrpG in all *hrpG\** samples, thereby validating its use for the analysis of HrpG regulons.

**Figure 2.**
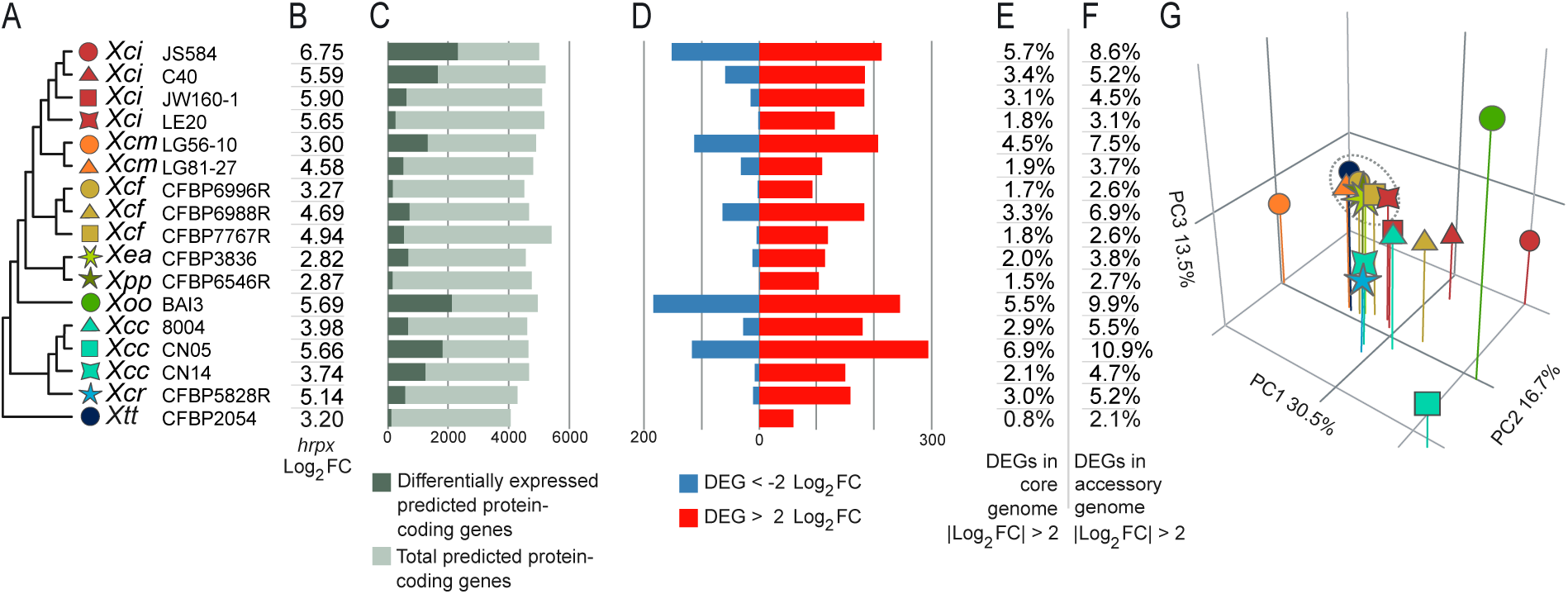
Features of the HrpG* regulons in 17 *Xanthomonas* strains. **A:** Phylogenetic tree of all strains as shown in Fig. 1A. **B:** Log_2_FC of *hrpX* in *hrpG\** strains compared to wild-type. **C:** Proportion of differentially-expressed predicted protein-coding genes per genome (AdjPval < 0.05). **D:** Number of DEGs (|Log_2_FC| > 2). **E:** Percentage of DEGs (|Log_2_FC| > 2) in the core genome only considering protein-coding genes. **F:** Percentage of DEGs (|Log_2_FC| > 2) in the accessory genome only considering protein-coding genes. **G:** Principal components analysis plot showing the first three principal components based on the average Log_2_FC of genes within each orthogroups of the core genome.

The number of differentially-expressed genes (DEGs, AdjPval < 0.05) varied across species from 137 to 2,355 genes (Fig. 2C; Table S2A). Considering a threshold of |Log_2_FC| > 2, the number of DEGs ranged from 59 to 429 (Fig. 2D). Notably, in all strains but *Xcf*_CFBP7767_ and *Xcf*_CFBP6996R_, the proportion of HrpG* regulated protein-coding genes in the core genome was significantly lower than in the accessory genome (Chi-square, BH FDR, AdjPval < 0.05, Fig. 2EF). Furthermore, the regulation of orthogroups by HrpG* within the core genome was highly variable, as highlighted by a PCA (Fig. 2G). In addition, the strains showed substantial differences in enriched Gene Ontology (GO) terms amongst DEGs considering a threshold of |Log_2_FC| > 2 (Fig. 3). Notably, only GO terms associated with protein secretion and protein secretion by the type III secretion system were enriched in the regulons of all strains, illustrating that the functions of the accessory HrpG regulon are highly diverse (Fig. 3).

**Figure 3.**
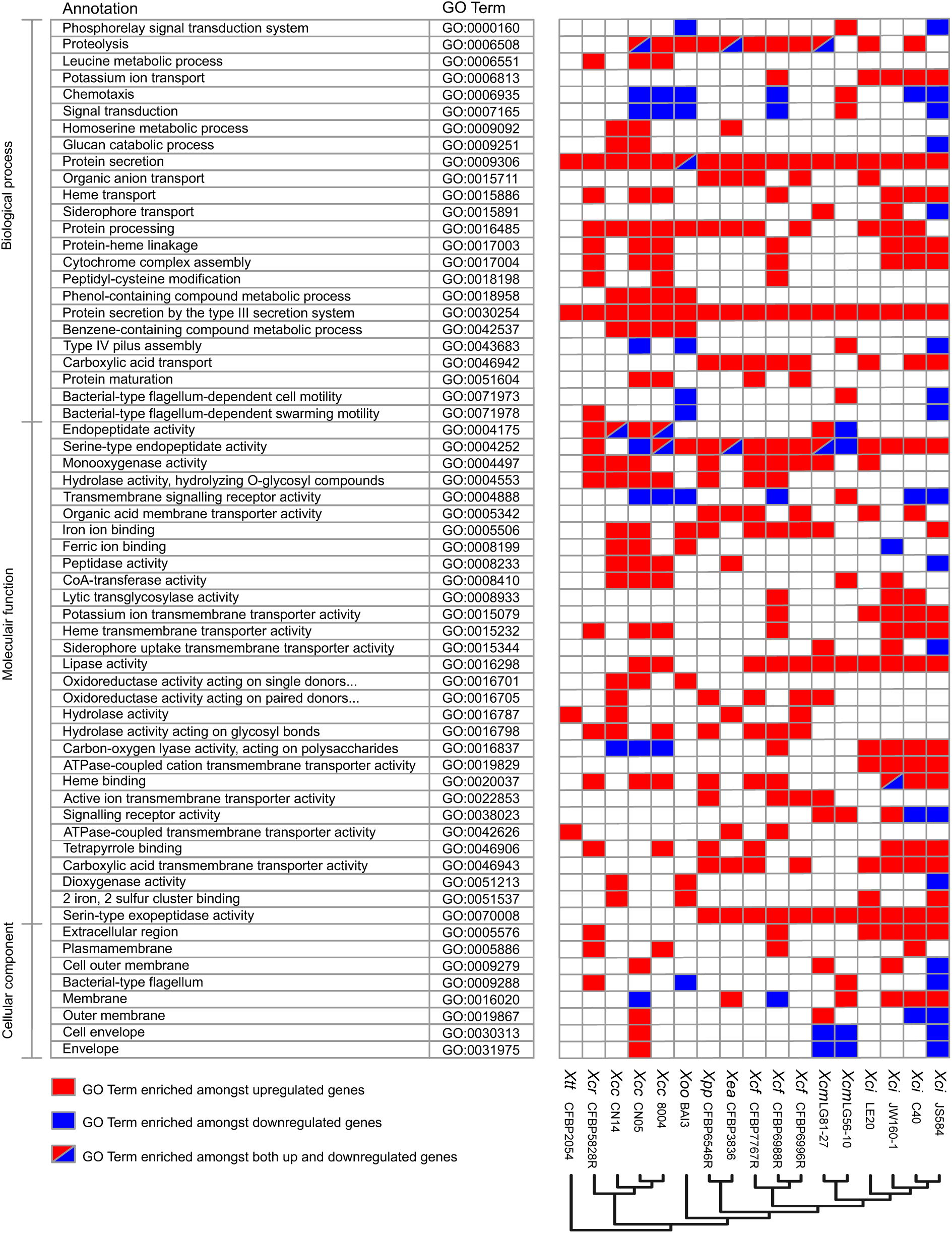
GO term enrichment in the HrpG* regulons in 17 *Xanthomonas* strains. Only GO terms enriched amongst regulated genes (|Log_2_FC| > 2) identified in at least three strains are depicted in the table to highlight the main commonalities across the strains (62 GO terms). A total of 197 unique GO terms were found to be enriched in at least one strain as detailed (Table S2B).

### The HrpG* core regulon is limited to only 26 orthogroups

To determine the HrpG* core regulon, orthogroups of the core genome of which at least one ortholog was differentially regulated in *hrpG\** samples across all strains were identified (Table 1). The identified HrpG* core regulon comprises one orthogroup coding for HrpX, 21 orthogroups encoding structural components of the T3SS and two orthogroups coding for the T3Es XopR and XopL. Additionally, the core regulon also includes two orthogroups coding for a putatively Sec/SPI-secreted acid phosphatase of the PAP2 superfamily (annotated as PhoC) and a major facilitator superfamily (MFS) transporter. Here, *hrpG* upregulation was ignored as sequence reads predominantly originated from plasmid-borne *hrpG*\* transcripts (Data S4 https://doi.org/10.57745/9OTYNJ). Thus, the identified HrpG* core regulon is almost exclusively associated with type III secretion.

**Table 1.** *Xanthomonas* HrpG* core regulon members.

| Gene annotation | Orthogroup number | Average log<sub>2</sub>FC<sup>a</sup> | SD Average log<sub>2</sub>FC |
| --- | --- | --- | --- |
| **Type III secretion system** |  |  |  |
| *hpaH/hpa2* | OG0000337 | 6.8 | 1.7 |
| *hrcV* | OG0000338 | 4.7 | 1.8 |
| *hrcQ* | OG0000339 | 5.6 | 2 |
| *hrcC* | OG0001932 | 6.6 | 1.5 |
| *hrcT/hrpB8* | OG0001933 | 5.1 | 1.4 |
| *hrpB7* | OG0001934 | 5.2 | 1.5 |
| *hrcN/hrpB6* | OG0001935 | 5.4 | 1.5 |
| *hrcL/hrpB5* | OG0001936 | 6.1 | 1.6 |
| *hrpB4* | OG0001937 | 5.6 | 1.5 |
| *hrpB3* | OG0001938 | 6.2 | 1.7 |
| *hrpB2* | OG0001939 | 6.7 | 1.4 |
| *hrpB1/hrpK* | OG0001940 | 7 | 1.4 |
| *hrcU* | OG0001941 | 6.4 | 1.6 |
| *hpaP/hrpC3* | OG0001942 | 5.5 | 1.4 |
| *hrcR/hrpD2* | OG0001943 | 5.6 | 1.6 |
| *hrcS* | OG0001944 | 5.5 | 1.6 |
| *hpaA* | OG0001945 | 5.1 | 1.7 |
| *hrcD* | OG0001946 | 6 | 1.6 |
| *hrpD6* | OG0001947 | 5.9 | 1.5 |
| *hrpE* | OG0001948 | 6.2 | 1.5 |
| *hpaB* | OG0001949 | 5.8 | 1.9 |
| **Type III effectors** |  |  |  |
| *xopL* | OG0000114 | 4.5 | 1.4 |
| *xopR* | OG0000535 | 3.4 | 1.1 |
| **Others** |  |  |  |
| *phoC* | OG0000410 | 3.3 | 1.6 |
| MFS transporter | OG0001668 | 3.5 | 1.5 |
| *hrpX* | OG0001966 | 4.6 | 1.2 |

<sup>a</sup> Of all shown orthogroups, at least one gene copy was significantly differentially regulated in all investigated strains (AdjPval < 0.05).

There is compelling evidence that group-I *Xanthomonas* acquired the type III secretion system cluster independently of the group-II *Xanthomonas* species, potentially resulting in the existence of different HrpG regulons (23). As the majority of strains used in this study belong to group-II *Xanthomonas* species, their core regulon was also identified. The identified group-II core regulon includes an additional 12 orthogroups compared to the *Xanthomonas* core regulon (Table 2). These include orthogroups encoding several known virulence determinants, such as a chorismate mutase and LipA whose function is independent of the T3SS and represent conserved and ancestral virulence mechanisms (24).

**Table 2.**
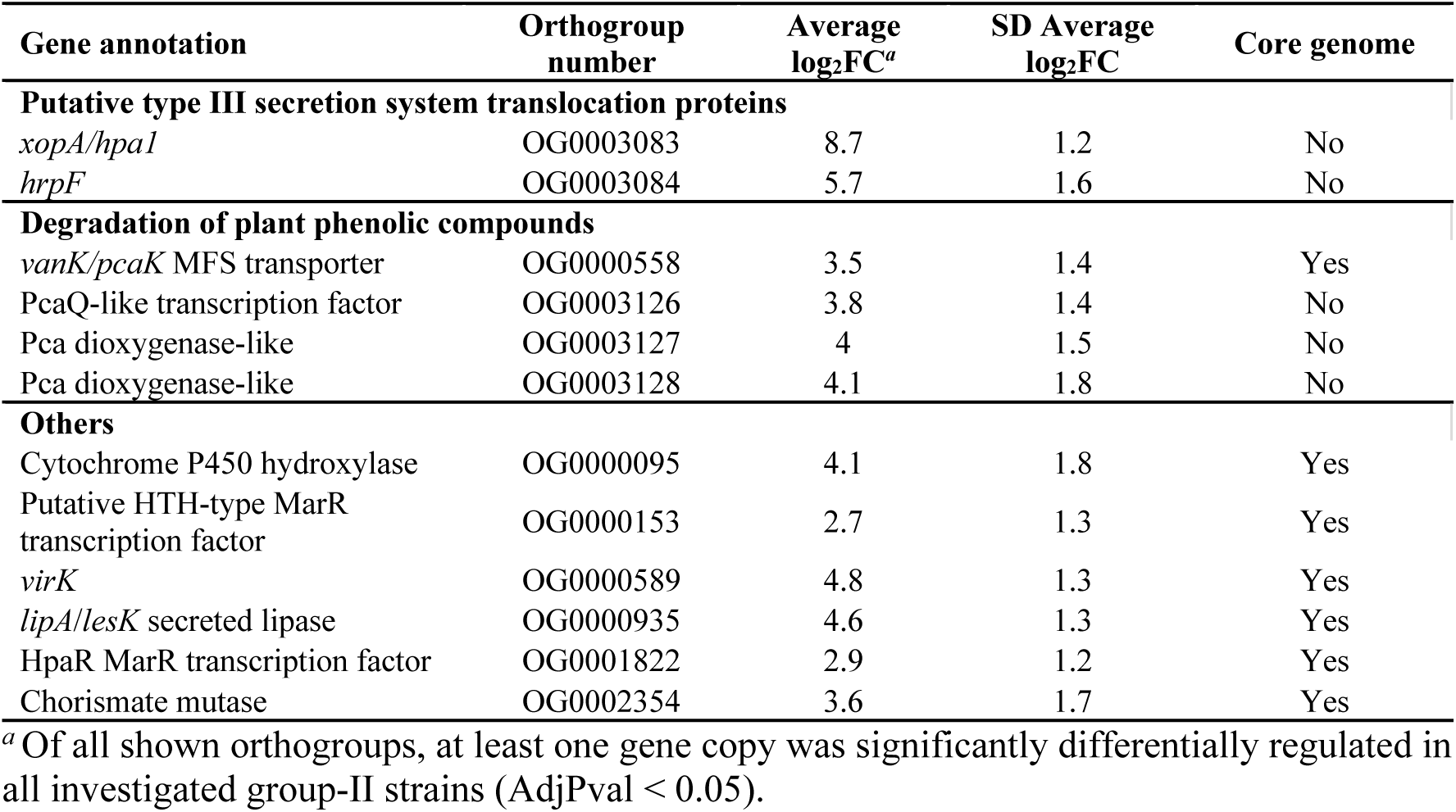
Group-II-specific *Xanthomonas* HrpG* core regulon identified in this study.

### Expression of most but not all T3Es encoding genes is controlled by HrpG*

Previous studies have shown that T3Es, both with and without a PIP-box motif in their promoters (TTCGB-N_15_-TTCGB), can be under positive regulation by HrpG or HrpX (3, 14–16, 20). To investigate the regulation of T3Es by HrpG* in the 17 strains analysed, we used the Effectidor software (25) to identify orthogroups encoding T3Es. A total of 72 orthogroups were predicted to contain orthologs coding for T3Es (Table S3A). From these, 12 orthogroups corresponding to *xopA*, *hrpW*, *hpaA* and other T3SS associated genes, which are generally not considered T3Es, were excluded (Table S3C). Additionally, Effectidor identified 15 singletons that are predicted to encode T3Es. Consistent with previous findings, our observations show that the majority of T3E genes are under the control of HrpG* (Table S3B; Fig. 4). However, the expression of few predicted effectors was not HrpG*-dependent, as observed for *xopAW*. As for the core T3E gene *xopM*, its expression was HrpG*-dependent except for both *Xcm* strains. Interestingly, most orthologs encoding transcription activator-like effectors (TALEs, OG000009) were positively regulated by HrpG*, although *Xci* TALEs were previously reported as expressed in a HrpG/HrpX-independent manner (14, 20). We thus conclude that the expression of most, but not all of the predicted T3E genes is under the positive control of HrpG*.

**Figure 4.**
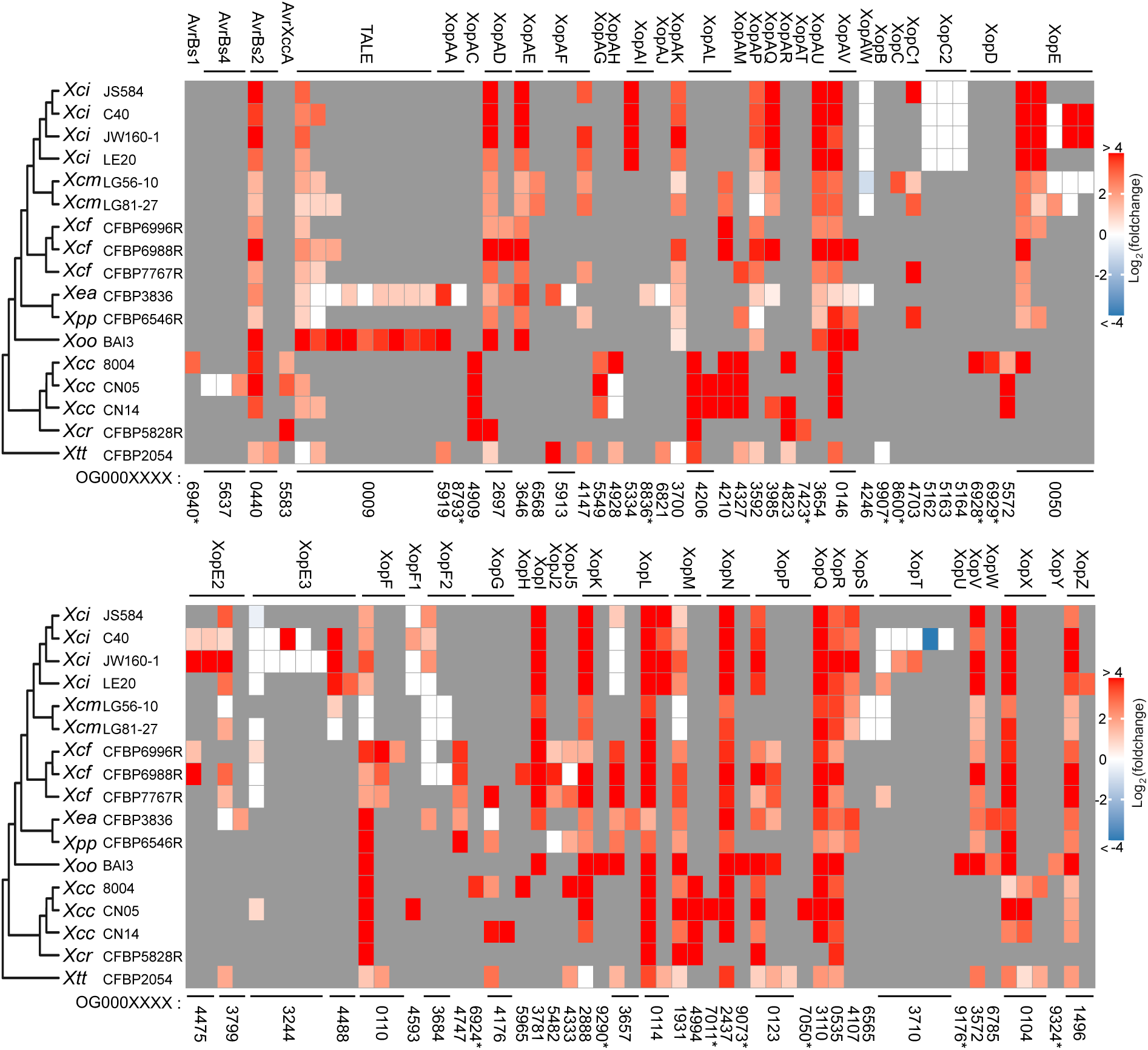
Regulation of orthogroups (OG) encoding putative T3Es in *Xanthomonas* strains expressing *hrpG\**. Only significantly DEGs (AdjPval < 0.05) are colored. Missing orthologs are marked in gray. Orthogroup annotation is given on top of the figure, orthogroup number at bottom of the figure. Asterisks indicate protein-coding genes which are singletons.

### Chemotaxis and motility are differentially regulated by HrpG* at both inter- and intra-specific levels

The PCA, based on the average HrpG*-dependent Log_2_FC of expression of orthogroups within the core genome, indicated that the regulatory network of HrpG* within the core genome differed considerably between strains, even for those of the same pathovar (Fig. 2G). Orthogroups that correlated strongly with the first three principal components were enriched for GO terms associated with the biological process of chemotaxis (Table S4). We therefore investigated the regulation of orthogroups comprising genes encoding methyl-accepting chemotaxis proteins (MCPs), Che signalling genes and structural components of the flagellum and the type-IV pilus. Interestingly, the regulation of these orthogroups was HrpG*-dependent in half the strains and both HrpG*-dependent upregulation and downregulation of these processes could be observed, depending on the strain (Fig. 5). For example, *Xcm*_LG56-10_ showed strong upregulation of genes in motility- and chemotaxis-related orthogroups while *Xcm*_LG81-27_ did not. These findings highlight diverse scenarios of HrpG*-dependent regulation of chemotaxis and motility in the *Xanthomonas* genus.

**Figure 5.**
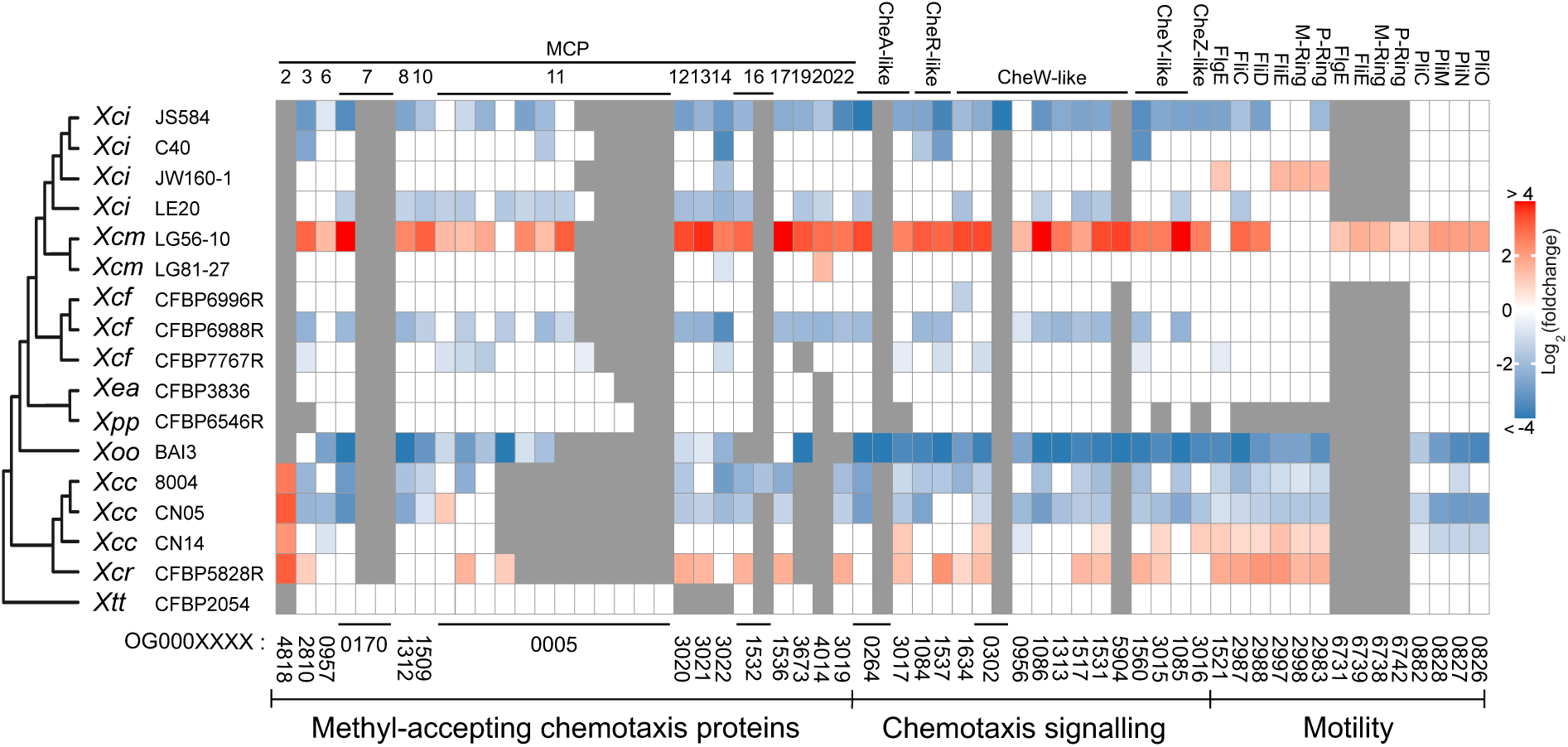
Regulation of orthogroups (OG) in chemotaxis and motility in *Xanthomonas* strains expressing *hrpG\**. Only significantly DEGs (AdjPval < 0.05) are colored. Missing orthologs are marked in gray. Orthogroup annotation is given on top of the figure, orthogroup number at bottom of the figure. Due to the extensive number of orthogroups potentially involved in chemotaxis, only those orthogroups that are regulated by HrpG* in at least two strains are shown, except for the secondary flagellar cluster specific to *Xcm* strains. Additionally, for both the flagellar and pilus gene clusters, only the regulation of key representative genes that encode various structural components of the flagella and pili are shown. MCPs: methyl-accepting chemotaxis proteins.

### Strain-dependent variability in HrpG*-mediated regulation of T2SS substrates

The HrpG* regulon of different strains exhibited significant enrichment in genes linked to GO terms associated with carbohydrate-active enzymatic activity and proteolytic activity (Fig. 3). Such genes play a crucial role in *Xanthomonas* virulence due to their involvement in the degradation of plant cell wall components, which facilitates nutrient acquisition and the efficient translocation of T3Es (19, 26). Typically, proteins encoded by these genes are characterized by the presence of a Sec/SPI signal peptide, indicative of secretion by the T2SS. Indeed, such signal peptides were predicted in 546 of the 3351 protein-coding genes of the pan HrpG* regulon, representing a significant enrichment for the presence of a signal peptide amongst regulated genes (|Log_2_FC| > 2, Chi-square, p < 0.0001, Table S2C). We therefore hypothesized that HrpG* might also regulate the T2SS, as previously shown for *Xanthomonas euvesicatoria* pv. *euvesicatoria* (*Xev*) (19). However, genes encoding the T2SS were not under the control of HrpG* (|Log_2_FC| > 2), except for *Xcm*_LG56-10_ (Fig. S1).

Though HrpG is known to regulate the expression of genes encoding proteins with carbohydrate active enzymatic and/or proteolytic activity (15, 17–19, 27, 28), such regulation can either be positive or negative without any obvious pattern (29). We thus examined the regulation of those genes in the transcriptomes of the 17 *Xanthomonas* strains (Fig. 6 and Fig. 7). Once more, various HrpG*-dependent regulatory patterns were observed. For example, orthologs in OG0003431, encoding putative S53 family serine proteases, were generally upregulated except in *Xoo*_BAI3_. Conversely, expression of orthologs in OG0000136, encoding putative S1 family serine proteases, were mainly downregulated, while *Xoo*_BAI3_ strongly upregulated one specific ortholog. Genes encoding proteins with putative pectin-lyase activity were also variably regulated by HrpG*. For example, orthologs in OG0000700, corresponding to genes encoding putative secreted GH28 family pectin lyases, were strongly upregulated in all strains except *Xoo*_BAI3_ and *Xtt*_CFBP2054_. In contrast, orthologs in OG0000100 and OG0000105, corresponding to genes encoding putative secreted PL1 family pectin lyases, showed considerable downregulation, especially in *Xanthomonas campestris* pathovars, but not in others such as *Xpp*_CFBP6545R_. These results are in line with previous reports, highlighting the differential regulation of genes encoding proteins with specific carbohydrate active enzymatic and/or proteolytic activity by HrpG* across different *Xanthomonas* strains. Nonetheless, most orthogroups showed consistent HrpG*-dependent regulation across the majority of strains, evidencing the existence of conserved HrpG*-mediated regulatory patterns for genes encoding specific families of carbohydrate active enzymes and/or proteases in the *Xanthomonas* genus.

**Figure 6.**
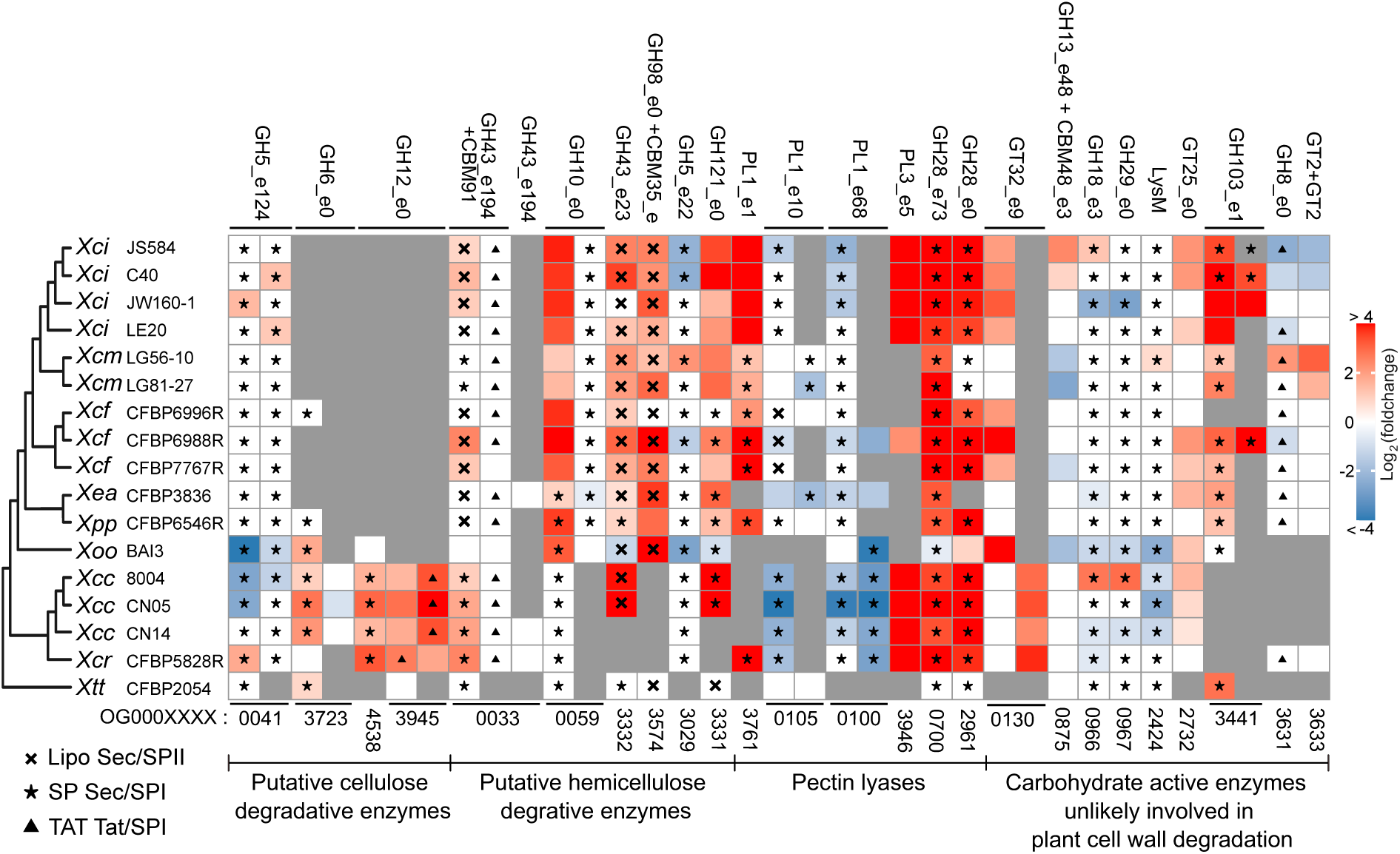
Regulation of orthogroups (OG) encoding carbohydrate active enzymes in *Xanthomonas* strains expressing *hrpG\**. Only genes with an AdjPval < 0.05 are colored. Absent orthologs are marked in gray. Orthogroup annotations are given on top of the panel according to their CAZy family. Orthogroup numbers are given at the bottom of the panel. Due to the extensive number of orthogroups encoding carbohydrate active enzymes, only orthogroups regulated in at least two strains are shown. The cross, star and triangle symbols indicate the presence of a signal peptide within the protein, as predicted by SignalP6.

**Figure 7.**
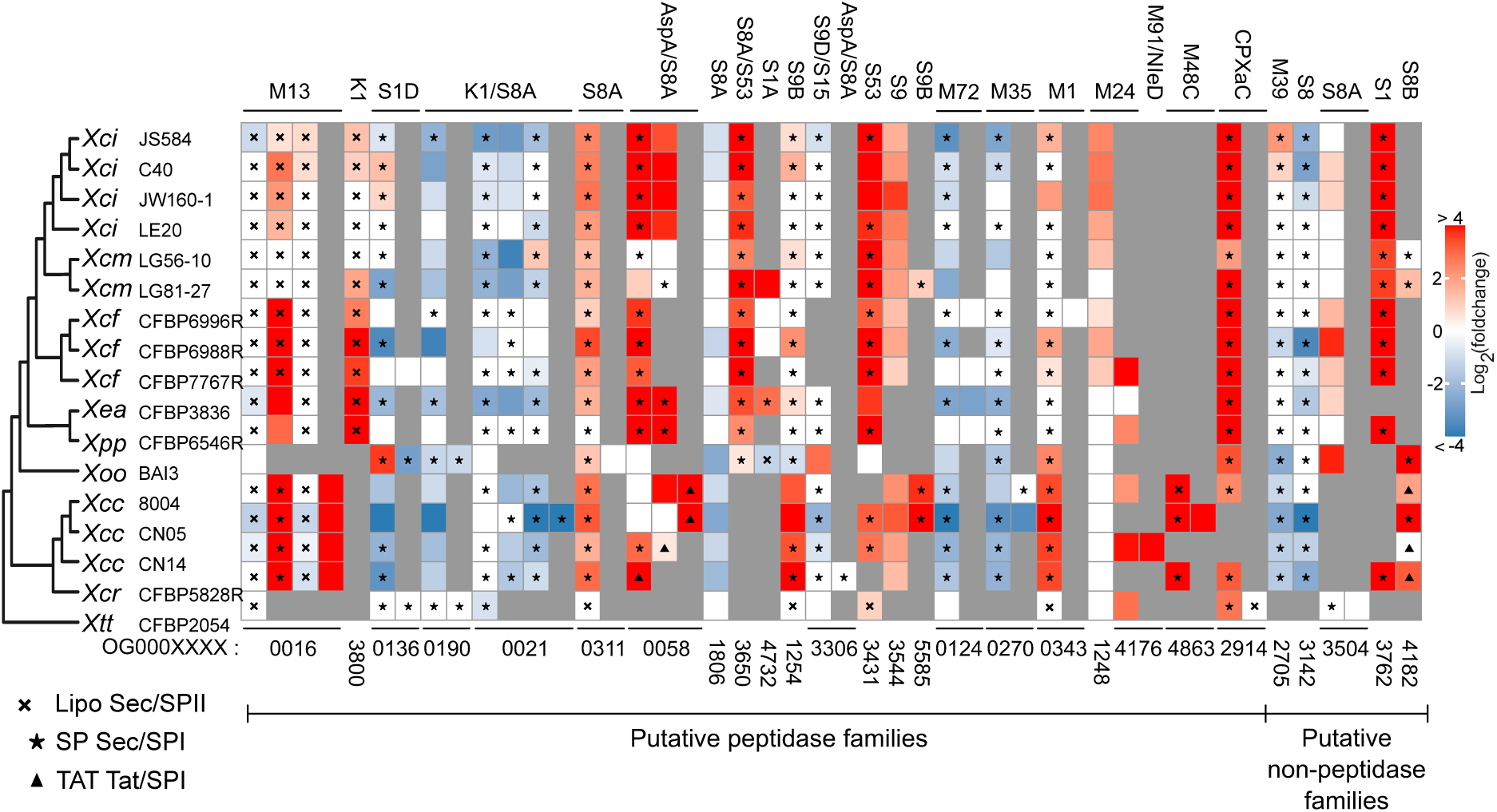
Regulation of orthogroups (OG) encoding proteins with proteolytic activity in *Xanthomonas* strains expressing *hrpG\**. Only genes with an AdjPval < 0.05 are colored. Absent orthologs are marked in gray. Orthogroup annotation is given on top of the figure, orthogroup number is given at the bottom of the figure. Orthogroups are annotated according to their MEROPS identifiers. Due to the extensive number of orthogroups encoding proteins with proteolytic activity, only orthogroups regulated in at least two strains are shown. The cross, star and triangle symbols indicate the presence of a signal peptide within a gene, as predicted by SignalP6.

### The HrpG* regulon members prepare *Xanthomonas* cells for the degradation of plant-derived phenolic compounds

The degradation of plant cell walls mediated by T2SS substrates in *Xanthomonas* species leads to the release of diverse phenolic compounds, including hydroxycinnamic acids, vanillic acid and 4-hydroxybenzoic acid (4-HBA). Interestingly, the HrpG* core regulon comprises an MFS transporter (OG0001168) involved in 4-HBA uptake in *Xcc*_8004_ (Table 1) (30, 31). In addition, the group-II *Xanthomonas* regulon comprises orthogroups coding different proteins involved in the uptake (VanK/PcaK, OG0000558) and putative degradation (two subunits of a PCA dioxygenase, OG0001327, OG0001328) of phenolic compounds (30, 31) as well as a PcaQ-like transcriptional regulator (OG0001326), which regulates phenolic compound metabolism in other plant-associated bacteria (Table 2) (32, 33). Therefore, a broader survey of orthogroups relevant for phenolic compound metabolism was conducted (Fig. 8). We observed that expression of numerous genes involved in the import and degradation of such compounds was upregulated, while some genes putatively involved in their efflux were downregulated. Notably, many orthogroups involved in these processes were absent from the *Xtt*_CFBP2054_ genome. Collectively, these results suggest that HrpG* prepares the group-II *Xanthomonas* metabolism for the import and degradation of plant-derived phenolic compounds.

**Figure 8.**
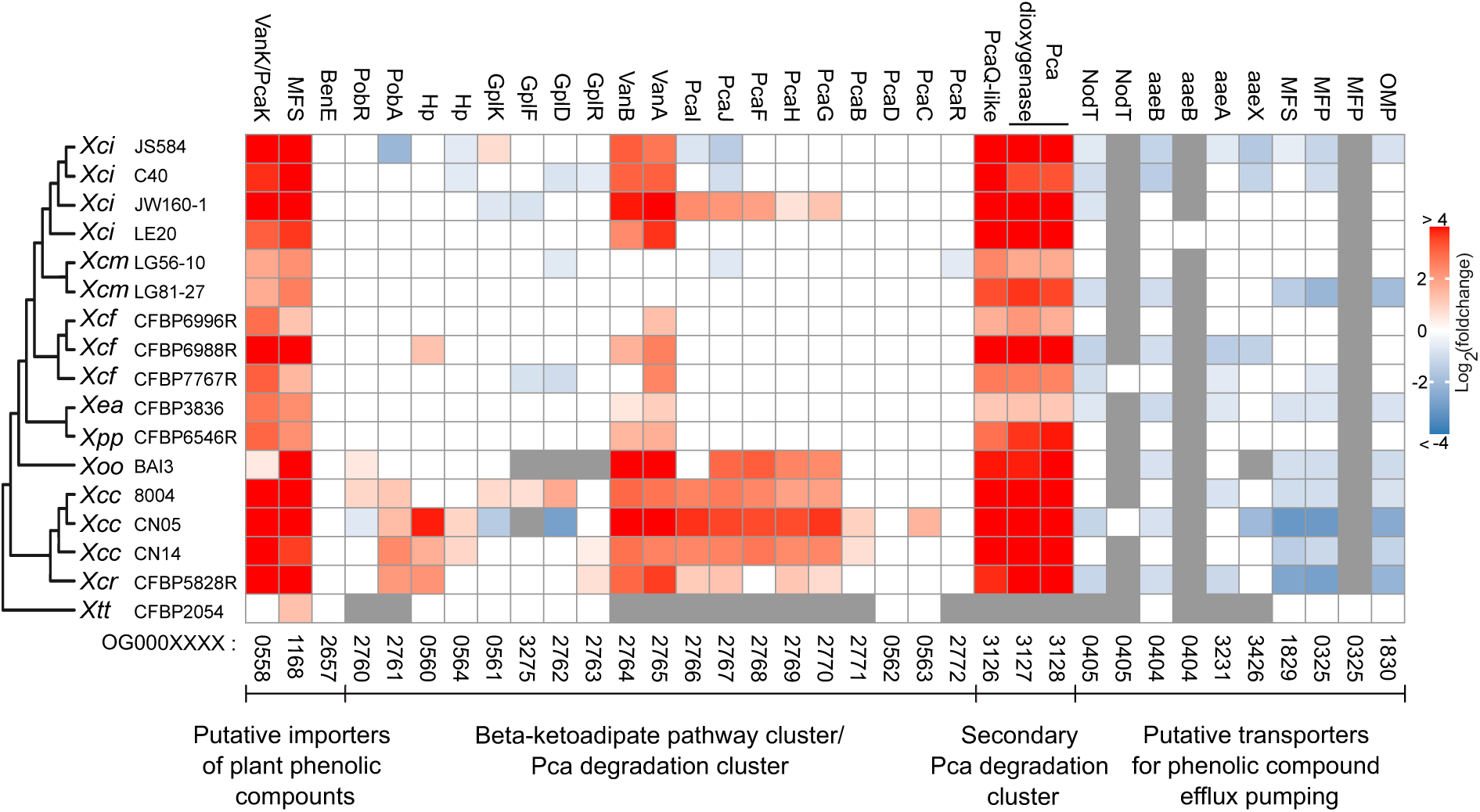
Differential expression of orthogroups (OG) encoding proteins putatively involved in plant phenolic compounds metabolism. Only genes with an AdjPval < 0.05 are colored. Absent orthologs are marked in gray. Orthogroup annotation is given on top of the panel, orthogroup number is given at the bottom of the figure. Pca: protocatechuate. Hp: hypothetical protein

### HrpG* induces the expression of genes involved in cytochrome C maturation in most *Xanthomonas* strains

GO terms associated with “iron binding”, “heme transport” and “siderophore uptake” were significantly enriched in the HrpG* regulons of different strains (Fig. 3) and are commonly associated with iron homeostasis. Orthologs of TonB-dependent receptors involved in iron uptake in *Xcc* (34) were differentially regulated in some strains expressing HrpG* (Fig. S2) but did not explain the observed enrichment of those three GO terms in all strains. Upon further investigation we found that the enrichment in these GO terms originated mainly from orthogroups comprising genes putatively involved in cytochrome C maturation (*Ccm* genes). Cytochrome C maturation complexes can have multiple roles in bacteria, including respiration (35), resistance to antimicrobial phenazines(36), or virulence as shown for *Xcc* (37). Interestingly, the identified regulated *Ccm* genes were located in an evolutionary-conserved gene cluster composed of a hypothetical protein, the sigma-factor RpoE4, another hypothetical protein with a zinc-finger domain and a S8A protease with a predicted Sec/SPI secretion signal (Fig. 9). Most genes in this cluster were regulated positively by HrpG* except in *Xpp*_CFBP6546R_ and *Xtt*_CFBP2054_. These results indicate that HrpG* regulates cytochrome C maturation in most *Xanthomonas* strains.

**Figure 9.**
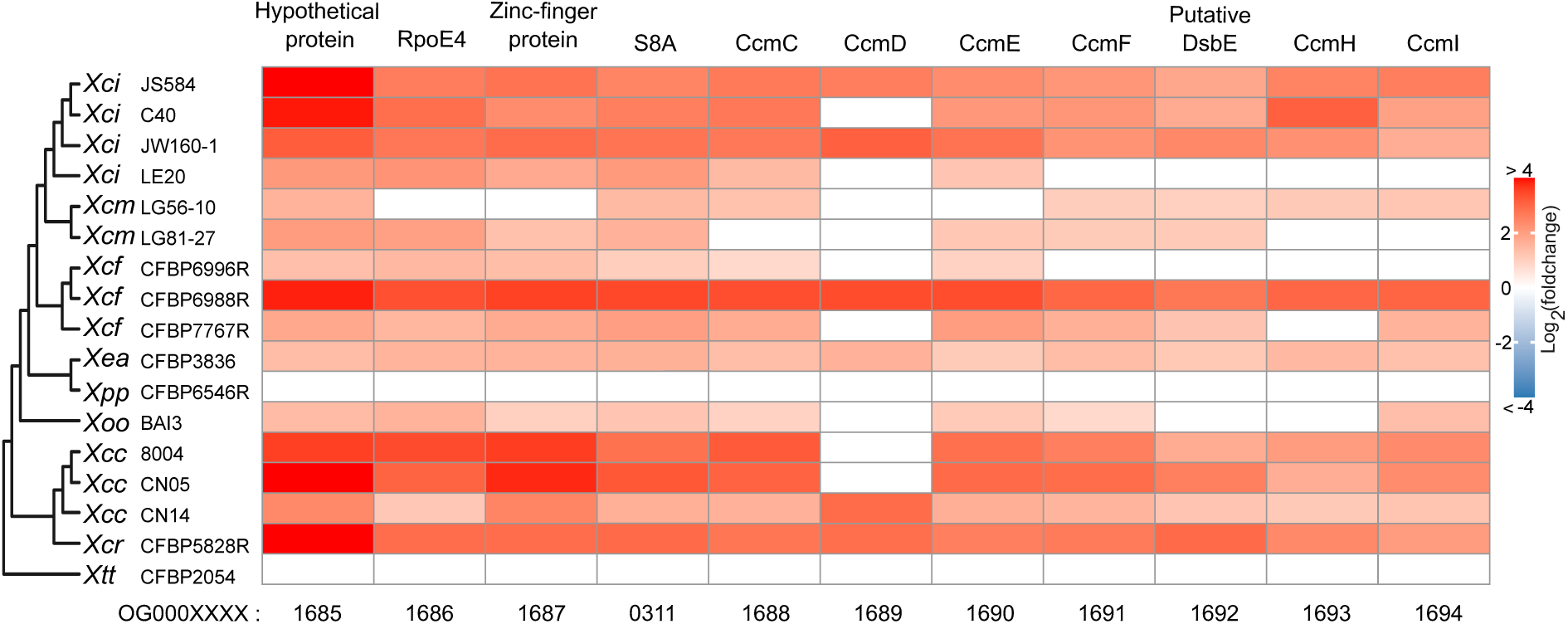
Differential expression of orthogroups (OG) encoding proteins putatively involved in cytochrome C maturation. Only genes with an AdjPval < 0.05 are colored. Orthogroup annotation is given on top of the figure. Orthogroup number is given at the bottom of the figure.

## DISCUSSION

This comparative transcriptomics study has provided a genus-wide overview of the evolutionarily-conserved processes regulated by HrpG*, as well as a glimpse into the intra- and inter-specific diversity of the regulon (Fig. 10). The regulons of the strains investigated here encompass hundreds, if not thousands of genes, including a variety of known virulence and adaptive pathways with notable enrichment for putative T2SS substrates. However, these regulons differ considerably, both within and between species. This variation is illustrated by a small HrpG* core regulon (26 orthogroups for the *Xanthomonas* genus and 38 for group-II *Xanthomonas*) and by the fact that only two gene ontology terms (GO0009306: protein secretion, and GO0030254: protein secretion by type III secretion system), were significantly enriched among the regulated genes across all strains. Additionally, our analysis revealed that the proportion of regulated protein-coding genes in the accessory genomes of most strains was significantly higher than that in the core genome. Collectively, these findings illustrate that the HrpG regulon from one *Xanthomonas* strain cannot be inferred from another related *Xanthomonas* strain. The diversity of HrpG regulon could be the results of long-term evolution process. Indeed, HrpG and HrpX were acquired prior to the split of *Xanthomonas* in group I and II and the acquisition of the T3SS (38) suggestive of an ancestral role in the regulation of other biological functions. How the T3SS became part of the core regulon of HrpG remains to be elucidated.

**Figure 10.**
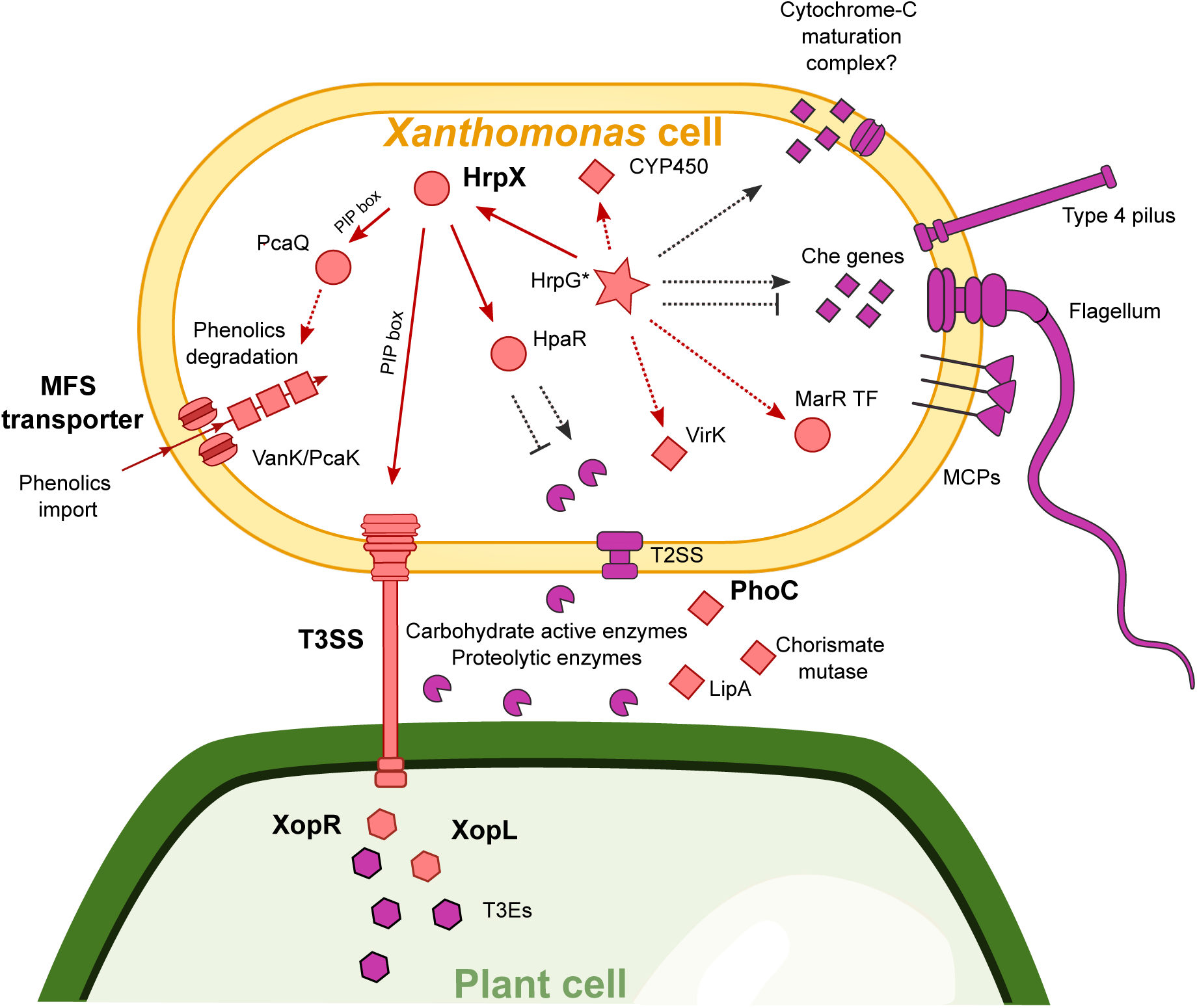
Schematic representation of the key processes identified to be regulated by HrpG* in this study. Red symbols annotated in bold indicate processes that are part of the *Xanthomonas* core regulon. Other red symbols indicate terms part of the group-II *Xanthomonas* core regulon. Symbols in purple highlight other regulated orthogroups which have been discussed. Dashed lines indicate hypothesized interactions, whereas continuous lines indicate interactions which are thought to be direct. T3SS: type III secretion system; T3Es: type III effectors; T2SS: Xps type II secretion system; MCPs: methyl-accepting chemotaxis proteins.

### The acid phosphatase PhoC belongs to the *Xanthomonas* HrpG* core regulon

The acid phosphatase PhoC (OG0000410) from the PAP2 superfamily belongs to the HrpG* core regulon and exhibits high conservation across *Xanthomonas* species. Of the 657 publicly available *Xanthomonas* genomes that contain a *hrp* T3SS gene cluster (Data S6, https://doi.org/10.57745/9OTYNJ), 650 harbour a *phoC* ortholog. Among these, 606 orthologs have a predicted Sec/SPI secretion signal whilst all others have a Sec/SPII secretion signal, indicating a strong conservation of T2SS-mediated secretion (Table S5A). The ortholog of *phoC* in *Xev* has been shown to be upregulated during the interaction with tomato although knockout mutants were not affected in virulence (39). Additionally, the *Xcc phoC* ortholog was not important for fitness inside cauliflower hydathodes nor for *Xcc* virulence on cabbage (40, 41). Despite the strong conservation of this orthogroup, its molecular or enzymatic functions remain elusive. In contrast, the functions of the T3Es XopL and XopR, both member of the HrpG* core regulon, have extensively been studied (42). The *xopL* orthologs encode atypical E3 ubiquitin ligases which can contribute to *Xanthomonas* virulence in several distinct mechanistical ways, depending on the species (43–47). This, together with significant interspecific variation in amino acid sequence, *in planta* subcellular localizations and host specific cell death-inducing capability indicate that this ancestral effector has undergone significant diversification (47). In *Xoo,* XopR is an effector which localizes to the plasma membrane, where it associates with various receptor-like cytoplasmic kinases (48–50). In both *Xoo* and *Xanthomonas axonopodis* pv. *manihotis* XopR is thought to contribute to virulence by interfering with PTI (48, 51).

### The group-II *Xanthomonas* HrpG* core regulon mediates ancestral T3SS-independent virulence mechanisms

The regulation of 12 orthogroups was identified to be HrpG*-dependent specifically in the group-II *Xanthomonas* species tested in this study. Among them, orthogroups encoding VirK, a chorismate mutase, and the LipA/LesK lipase have been previously identified by comparative genomic analyses to be potential conserved virulence determinants in *Xanthomonas* species (24). Orthologs of *virK* are conserved across various lineages of plant-associated bacteria and are present in the genomes of all 657 *Xanthomonas* strains with a T3SS (Data S6, https://doi.org/10.57745/9OTYNJ). While the function of VirK has not yet been experimentally addressed, *in silico* analyses suggest that it interacts with structural components of both the T3SS and the Xps T2SS, and it is hypothesized to be secreted by the T2SS (24). Orthologs of the chorismate mutase are also found in all genomes of *Xanthomonas* species with a T3SS, showing high levels of amino acid sequence similarity, including a Sec/SPI/SPII signal secretion signal (Table S5B; Data S6, https://doi.org/10.57745/9OTYNJ). Although its molecular functions remain undefined, it contributes to *Xoo* virulence, possibly by interfering with salicylate-dependent immunity (24, 52) (Degrassi *et al.*, 2010; Assis *et al.*, 2017). LipA/LesK orthologs are conserved with a Sec/SPI secretion signal in nearly all *Xanthomonas* species with a T3SS but absent from non-pathogenic *Xanthomonas* (Table S5C; Data S6, https://doi.org/10.57745/9OTYNJ) (24). Interestingly, while an ortholog is present in *Xtt*_CFBP2054_ genome, it lacks a PIP-box promoter motif, likely explaining the HrpG*-independent expression of *LipA/LesK* in this Group-I *Xanthomonas*. *Xev, Xci* and *Xoo* LipA/LesK contribute to virulence on tomato (53), citrus (24) and rice (54), respectively. Importantly, the ortholog of LipA/LesK is also a virulence mechanism for *Xylella fastidiosa* on grapevine (55) suggestive of an ancestral virulence function in the genera *Xanthomonas*, *Xylella* and *Burkholderia* (24) before its co-optation in the HrpG regulon of Group-II *Xanthomonas*.

### The group-II *Xanthomonas* HrpG* regulon relies on a transcriptional regulatory cascade

In addition to HrpX, the group-II core regulon comprises three additional transcriptional regulators, namely a MarR transcription factor, HpaR and a PcaQ-like transcriptional activator. The *Xci* MarR ortholog is essential for pathogenicity on Rangpur lime (56) although its gene targets are unknown. The transcriptional regulator HpaR is essential for *Xcc* virulence and thought to regulate the expression of various virulence mechanisms including extracellular proteases secreted by the Xps T2SS (17). As for the PcaQ-like transcriptional activator, it is hypothesized to be the main regulator of genes involved in protocatechuate degradation pathway, which is known to be important for *Xcc* virulence (30, 31). Interestingly, in *Xtt*_CFBP2054_, the PcaQ-like transcriptional activator is absent and the expression of the MarR transcription factor and *hpaR* is HrpG*-independent. This could explain the great divergence observed between the regulons in *Xtt*_CFBP2054_ and those in group-II *Xanthomonas*. These results thus suggest that a complex cascade defines the full extent of HrpG* regulons in group-II *Xanthomonas*, which could be refined by determining the transcriptomes of the corresponding single and multiple mutants in a *hrpG*\* mutant background.

### HrpG* regulates motility and chemotaxis in unpredictable ways

Motility and chemotaxis are known virulence determinants in *Xanthomonas*, primarily associated with the initial stages of infection (57–60). As HrpG expression gradually increases during infection (22, 61), it seems likely that HrpG could mediate the suppression of motility and chemotaxis, thereby potentially limiting PTI induced by flagellar components (29). So far, reports in *Xci* have indeed shown that HrpG negatively regulates motility and chemotaxis, likely in a HrpX-independent manner (14). However, we report the variable regulation of these processes. For instance, HrpG*-dependent downregulation was observed in *Xoo*_BAI3_, while no regulation was observed in *Xcm*_LG81-27_ and, remarkably, upregulation was measured in *Xcm*_LG56-10_. These findings are intriguing, especially considering the crucial role of motility and chemotaxis in the virulence of *Xanthomonas* species, and suggest the presence of complex inter- and intra-specific virulence mechanisms that require further investigation. In conclusion, we demonstrate that *Xanthomonas* species not only differ by their gene contents (Fig. 1), but also by their gene expression profiles, which are diverse even at the intra-pathovar scale. These observations are consistent with a general evolutionary context where virulence factors important in some hosts are often directly or indirectly recognised as immune elicitors in other hosts and therefore undergo diversifying selection. The observed differential expression within the HrpG regulon may limit recognition of some of these genes while diversifying the strains encountered by plants. Such adaptative transcriptional regulation would have the advantage of being transient, plastic and environment-dependent thus facilitating the emergence of novel virulence properties.

## MATERIALS AND METHODS

### Bacterial strains, plasmids and growth conditions

Bacterial strains and plasmids used in this study are listed (Table S1). *Xanthomonas* strains were grown at 28 °C in MOKA medium (34). *Escherichia coli* cells were grown on LB medium at 37 °C. For solid media, agar was added at a final concentration of 1.5 % (w/v). Antibiotics were used when appropriate at the following concentrations: 50 μg/mL kanamycin, 50 μg/mL rifampicin, 40 μg/mL spectinomycin.

### Genome sequencing and assembly

Genomic DNA was extracted from bacterial cells grown overnight in MOKA-rich medium using the Wizard genomic DNA purification kit (Promega) or as described (62) for short- or long-read sequencing, respectively. Shotgun sequencing of genomic DNA was performed either on HiSeq2000 Illumina platform (63) or PacBio (64). Genome assembly was performed as described for short read assemblies (63). FLYE Assembler (version 2.9) was used for long-read assemblies (65). The GenBank accession numbers for the genomes generated in this study are given in the Table S1.

### Cloning of pBBR-hrpG* plasmid

The *hrpG*\* (E44K) coding sequences were amplified from *Xcc* strain 8004 *hrpG*\* (66) and *Xanthomonas vesicatoria* pv. *euvesicatoria* strain 85-10 *hrpG* (1) and cloned into pBBR1-MCS-2 (67) as described (16) (Table S1D). Plasmids were introduced into *E. coli* by electroporation. The plasmids were introduced into the different *Xanthomonas* strains by triparental mating using pRK2073 as helper plasmid (68, 69) (Table S1D).

### RNA extraction, rRNA and tRNA depletion and cDNA pyro-sequencing

RNAs were extracted from *Xanthomonas* strains grown at exponential phase (OD_600 nm_ between 0.5 and 0.7) in MOKA rich medium. For each strain, RNA was extracted from at least three independent replicates of both the wild-type or empty vector conjugates and the HrpG* conjugates as described (22). Specific *Xanthomonas* probes were employed to deplete rRNA and tRNA molecules (16) after which RNAs were fractionated into short (<200 nt) and long RNAs using Zymo Research RNA Clean & Concentrator TM-5 columns (Proteigene). Strand-specific RNA sequencing was performed on an Ilumina HiSeq2000 platform, as described (22). A detailed overview of all generated and used RNA-Seq libraries and their NCBI Sequence Read Archive (SRA) accessions is provided in Table S1E.

### Structural annotation of *Xanthomonas* genomes

For each strain, the different RNA-Seq libraries available were merged in order to experimentally support the structural annotation of the genomes using Eugene-PP (70). The used RNA-Seq libraries are detailed in Table S1. The annotations used and generated in this study are provided in Data S1 (https://doi.org/10.57745/9OTYNJ).

### Differential expression analyses of the RNA-Seq results

RNA-Seq reads were pseudo mapped onto the reference genomes using Salmon (version 1.4.0) with standard parameters (71). An overview of which RNA-Seq datasets were used is given in Table S1E. Overall mapping quality was assessed using multiQC (72). DEGs were identified using DESeq2 with Salmon’s count tables using standard parameters (73). To assess sample reproducibility, PCA and MA plots were generated using DESeq2’s plotPCA and plotMA functions. Volcano plots for each strain were made using EnhancedVolcano (74). The PCA on the expression of all orthogroups in the core genome across all strains was based on a matrix with a single average Log_2_FC value for all genes within an orthogroup. Genes with non-significant Log_2_FC were assigned a Log_2_FC of 0. Session information, appropriate raw data and relevant R scripts to reproduce the results are available in a code repository (Data S5, https://doi.org/10.57745/9OTYNJ).

### Identification of the core HrpG* regulon

An orthogroup database constructed using Orthofinder (version 2.4.0) with standard parameters (75) was used to compare the regulation of genes by HrpG* across different strains. The species trees in Figure 1 were made using Orthofinder’s STAG algorithm (76) and rooted using STRIDE (77). Orthogroups, for which across all strains, at least one ortholog was differentially regulated in the wild-type or empty vector strains compared to the *hrpG\** strains (AdjPval <0.05) were considered orthogroups part of the core regulon. To investigate the regulation of the endogenous *hrpG* by HrpG*, we investigated whether reads mapping to *hrpG* originated from the endogenous *hrpG* gene or the exogenous *hrpG\** gene by mapping RNA-Seq reads to both endogenous and HrpG* sequences. Reads that did not map to these two sequences or which mapped multiple times were then removed. The coverage at the polymorphic site of *hrpG/hrpG\** for the remaining reads was then assessed. A Chi-square test was used to test for equal proportions of HrpG*-dependent regulation of predicted protein-coding genes within the core and accessory genomes. Multiple testing correction was performed using the Benjamini–Hochberg procedure (78).

### Conservation of the core HrpG regulon across all publically available genomes

To investigate the conservation of orthogroups identified in this study across plant pathogenic *Xanthomonas* strains with a T3SS, an additional orthogroup database was built using publically available genomes of *Xanthomonas* species with a T3SS (Data S6, https://doi.org/10.57745/9OTYNJ). This database was built on available genomes having less than 206 contigs and a Busco score of over 94.9% (657 genomes in total representing 25 *Xanthomonas* species and at least 67 different pathovars, Data S6). The conservation of relevant orthogroups identified in this study was studied for each corresponding orthogroup in the orthology analysis constructed with the 657 genomes.

### Gene ontology and signal peptide prediction

Enrichment of specific GO terms in various gene lists was investigated using the TOPGO R package (79). Signal peptides were predicted by signalP6.0 (80). A Chi-square test was used to test for an enrichment of genes with Sec/SPI amongst regulated genes. When appropriate, promoter sequences were scanned for PIP-box motifs using the TTCGB-N_15_-TTCGB consensus (3) allowing for one mismatch.

### Regulation of type three effector gene expression

T3Es were predicted using Effectidor (25) using the advanced mode, which makes use of the genomes GFF3 files to more accurately predict T3Es. Each orthogroup of which at least one ortholog was annotated as a true T3E by Effectidor was considered a true T3E orthogroup. True T3E orthogroups were further manually annotated and curated by integrating BLASTP, paperblast (81) and the EUROXANTH database (42). Because genes encoding TALEs are often poorly assembled due to their repetitive sequences (82), only TALE orthologs for which the complete N-terminal domain upstream of the repeat region was present were considered.

### Regulation of genes relevant for motility, carbohydrate active enzymes and proteases

Orthogroups involved in motility were identified using the Eugene-PP genome annotations and manually curated. Orthogroups encoding proteins with carbohydrate active enzyme activity were predicted using dbCAN3 (83) and further curated and annotated using the Carbohydrate-Active enZYmes (CAZY) Database (84). Genes encoding enzymes with protease activity were identified using the Eugene-PP genome annotations and manually curated and annotated using the MEROPS database (85).

### Visualization of regulation against phylogeny

Several figures were generated in order to explore a potential relationship between strain phylogeny and the regulation of specific orthogroups under HrpG*. The HrpG*-dependent Log_2_FC of expression all genes within an orthogroup were plotted against the phylogenetic tree as made by Orthofinder. The Log_2_FC of genes for which either Log_2_FC or AdjPval were assigned NA values by DESeq were set to 0. Figures were built using GGtree with equal branch length for the species tree (86).

### Availability of data and materials

All genomes sequenced and assembled in this study have been made publicly accessible. For a comprehensive list of their DOIs, please see Table S1. The transcriptome sequencing reads are available in the NCBI Sequence Read Archive (SRA), with the specific SRA accession numbers provided in Table S1. Supplementary data files can be found on recherche.data.gouv.fr under the DOI: https://doi.org/10.57745/9OTYNJ. This includes generated genome annotation for the 17 strains as Data S1, and comparative statistics and orthogroup analyses for these genomes, performed using OrthoFinder, as Data S2. R scripts utilized for analyzing the data, are stored in a code repository in Data S5. Comparative statistics and orthogroup information for 675 publicly available *Xanthomonas* genomes which include a type T3SS are detailed in Data S6, along with their respective genome accession numbers.

## ACKNOWLEDGEMENTS

We wish to thank Claudine Zischek for the construction of *Xcc* strains, Aude Cerutti for helpful advice on ribodepletion, Caroline Bellenot for sharing curated *Xanthomonas* genomics resources and Naama Waagner for running Effectidor analyses.

## DATA AVAILABILITY

All genomes sequenced and assembled in this study have been made publicly accessible at bbric.toulouse.inra.fr. For a comprehensive list of their DOIs, please see Additional File 1. The transcriptome sequencing reads are available in the NCBI Sequence Read Archive (SRA), with the specific SRA accession numbers provided in Additional File 1. Supplementary data files can be found on data.gouv.fr under the DOI: https://doi.org/10.57745/9OTYNJ. This includes generated genome annotation for the 17 strains as Data S1, and comparative statistics and orthogroup analyses for these genomes, performed using OrthoFinder, as Data S2. R scripts utilized for analyzing the data, are stored in a code repository in Data S5. Comparative statistics and orthogroup information for 675 publicly available *Xanthomonas* genomes which include a type T3SS are detailed in Data S6, along with their respective genome accession numbers.

## AUTHORS CONTRIBUTION

The initial XANTHOMIX proposal was written under coordination of MA, OP, MAJ, LDN and RK. BR, SCU, BS, EC, AD, LDN, EL and LG were involved in the sequencing of four *Xanthomonas* genomes, the construction of the different *Xanthomonas* conjugates and the preparation and sequencing of cDNAs. SCA handled genome annotations, transcriptomic pseudomapping, and Orthofinder analyses. MFJ performed the initial transcriptomic analyses. TQM performed quality controls on the raw data, conducted the in-depth comparative transcriptomic analyses, prepared all the figures and drafted the manuscript. AB and LDN supervised the analyses, interpreted of the data and wrote the manuscript. All authors read, commented and approved the final manuscript.

